# Extracting human data from published figures: implications for data science and bioethics

**DOI:** 10.1101/376848

**Authors:** Brian J. Cox

## Abstract

The advent of text mining and natural text reading artificial intelligence has opened new research opportunities on the large collections of research publications available through journal and other resources. These systems have begun to identify novel connections or hypotheses due to an ability to read and extract information from more literature than a single individual could in their lifetime. Most research publications contain figures where data is represented in a graph. Modern publication guidelines are strongly encouraging publication of graphs where all data is displayed as apposed to summary figures such as bar charts. Figures are often encoded in a graphing language that is interpreted and displayed as a graphics. Conversion figures in publications to the underlying code should enable text-based mining to extract the underlying raw data of the graph. Here I show that data from publications greater than 15 years old that contain time series data on human patients is extractable from the original publication and can be reassessed using modern tools. This could benefit cases where data sets are not available due to file loss or corruption. This may also create and issue for the publication of human data as sharing of human data often requires research ethics approval.

**Author summary:** Figures embedded in published research manuscripts are a minable resource similar to text mining. Figures are text based code that draws the image, as such the underlying text of the code can be used to reassemble the original data set.

## Introduction

In the field of data science, online documents and publications are resources of data relating anything from corporate finance to human health and disease. Artificial intelligence based text readers use publications to glean information on a wide variety of topics and propose novel hypotheses(1). Most publications also contain graphical representations of data, which is an essential method to convey research findings in nearly all fields of study such as physics, biology or psychology. Often the raw data underlying most figures involving human subjects are not available where study subject consent, research ethics or institutional review boards often protect individualised data(2,3).

To gain access to subject data from publications involving human medical research applying to a IRB/REB for approval for de-identified data access is often not granted. In general research data on human subjects is not shared to the same extent as gene expression data. In all cases, data is anonymized when it is accessible, which is an essential restriction for human subject data. In publications, graphing of individual data points is encouraged by many journals to improve the informative representation of the data and research integrity, as opposed to compressed representations such as bar graphs, box plots or means(4,5). In many publications we can see the plots of the human patient data but cannot get access to the data for reanalysis using novel statistical methods or merging data sets.

Portable Document Format (pdf) based publications are a standard in publications. The images within PDF are encoded as Scalable Vector Graphics (SVG) a mark-up language. I reasoned that information about the data points was preserved in the SVG markup language that generates the data graphs. Key to this assumption is that the data was graphed in the order that it appeared in the data table used to generate the figure. Therefore reading the graphic’s code file should enable the recreation of the original data. The availability of this data information is double-edged, as it benefits a growing interest and needs for open source data sharing, experimental reproducibility and data integrity(6), but may violate the REB/IRB approval for use and publication of human subject data(2).

## Results and Discussion

I began by opening a PDF of a 2004 publication in the New England Journal of Medicine reporting the measurement of a serum protein at multiple time points during pregnancy comparing healthy and preeclamptic pregnancies(7). I have not contacted the authors to request the data used to generate the figure, I used this figure for illustrative purposes only.

The page with the figure of interest was extracted (Fig 1A). To simplify the data mining I removed text and graphical information, such as axis labels and tick marks that did not represent the desired data points (Fig 1B) and saved a scaled vector graphics file (Fig 1C, S2 file). For time-resolved data, such as consecutive draws of patient blood and measurement of serum protein concentrations, I noted that the data in the SVG file, represented as path objects, followed a repetitive pattern of increasing X coordinate values representing increasing time points of a patient. Then the X values lowered and repeated the increasing pattern, indicating that the next patient was being graphed. Next, I noted that the data in the original figure was represented as two different colours, which was encoded in the path object as a colour code. These differences were encoded into a text parser to identify path objects and score them into a table a series of patients and their status as either a healthy or preeclampsia case (S1 file). The path object’s code contained X and Y coordinate data that represented the objects physical placement on the original graph. Extracting the coordinate information created the values for gestational age at sampling and the corresponding serum protein concentration for that time point (Fig 1D). I determined the accuracy of the data table generated from the parsed SVG file by ensuring that the same numbers of patients were returned for each disease state (healthy and preeclampsia) as reported in the manuscript. While the data set is an accurate reproduction of the plot, the data is scaled to the Cartesian points, but can be easily rescaled to the original data ranges(compare Figs 1 A and E, S1 file).

**Fig 1.**
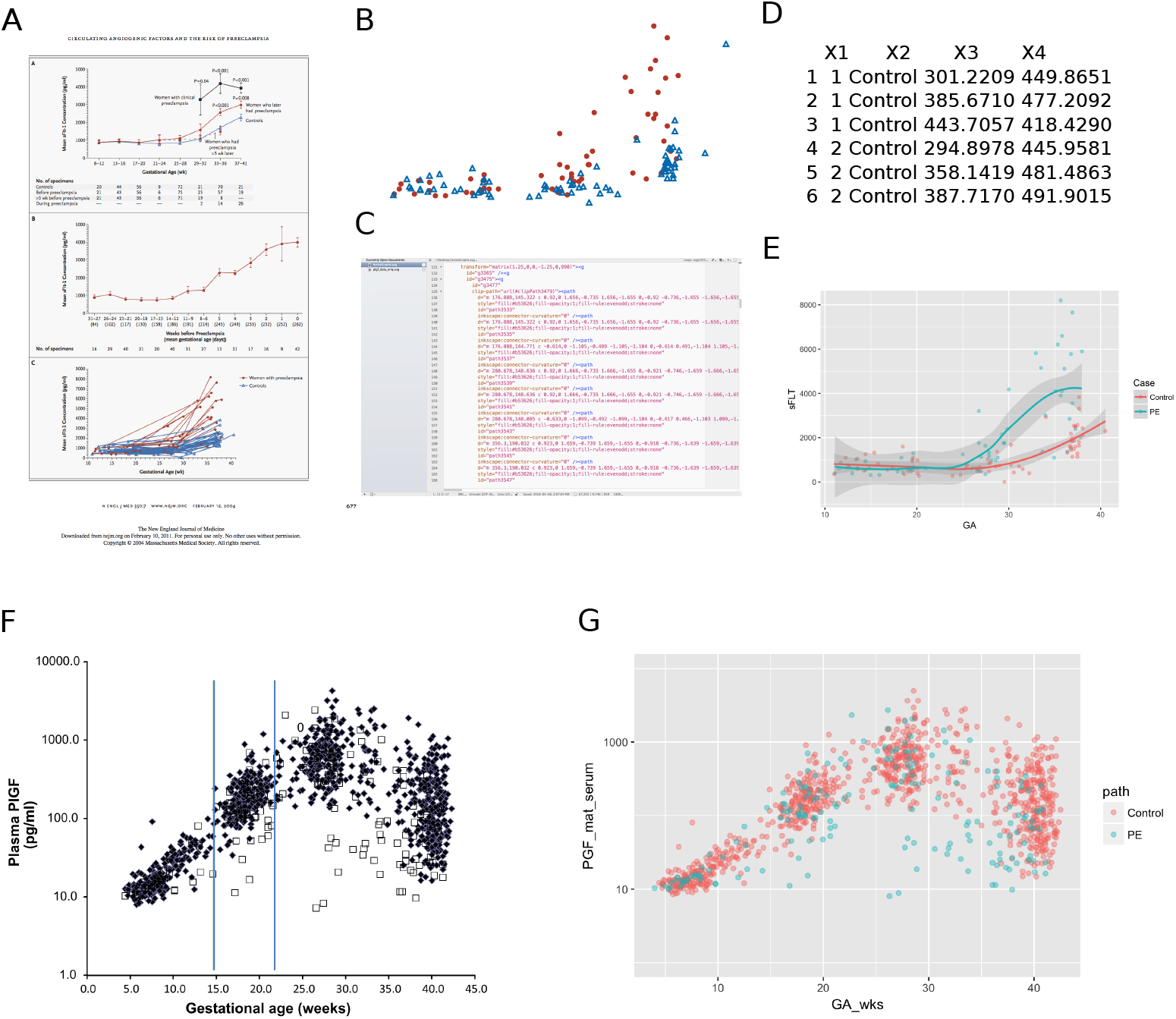
Workflow to extract and assess raw data from published figures. A) The page containing a figure with data of interest (lower figure panel). B) After cleaning only the data points are present. C) Example of the underlying code for the SVG file used to generate the figure in B. D) The extracted table containing patients (X1) as series with diagnosis (X2), but data is not scaled to gestation weeks (X3) or serum protein concentrations (X4). E) Final plot rescaled to match the published values and data range. F) a second example where patient classes are represented as shapes and data is log scaled. G) Final plot of extracted data.

In the second example PGF serum measurements during multiple time points in pregnancy were measured in healthy and preeclampsia patients(8) (Fig 1 F). The original figure was plotted as log scale, and I extracted and rescaled the data (Figs 1 F and G, S1 and S2 files). Additionally, in this example patients were graphed as squares and diamonds, the parser was modified to identify differences in path objects related to shape rather than colour, as in the first example. After parsing, I identified 299 patients while the article reported 300. Inspection of the graph and SVG file indicated that the original graphic had one patient case plotted as a “0” rather than a square or diamond shape that represented healthily and preeclampsia cases. This finding explained the failure of the parser to extract this data point.

Other errors were noted in two other extracted figures (data not shown). In these, either the number of samples plotted in the figure did not match the figure description or methods statements in the article (data not shown) or samples in a patient series became mislabelled (transposing from preeclampsia to healthy annotation). These graphing errors are more concerning, as it is unclear if these errors affected other data tables or calculations within these publications.

The implications of these findings for data science are considerable as all plotted data that is a scalable graphics file can be extracted this way. AI systems and automated code parsers could be developed to routinely or bulk extract the underlying data from published documents. Previous methods to extract information utilised image interpreters (https://automeris.io/WebPlotDigitizer). However, when large numbers of data points are over-plotted (plotted on top of each other), these methods fail to find all data points due to obscuring by features that overlap (data not shown). Furthermore, information on the order of the data points is lost by these image analysis methods (data not shown) but is preserved when parsed from the SVG code.

My findings have implications for research ethics, as the individual data was likely not intended to be published since these articles contained no associated tables of the patient data used to generate the graph. As a means of protecting individuals from purposeful or inadvertent identification, many REB/IRBs require human data to be published as group means or other aggregate statistics and not individual values. I reason that it is improbable for the anonymised data extracted from graphs to be able to re-identify individuals but may violate the original consenting of the patients or institutional policy. The free availability of this human subject data means that IRBs, REBs and governments may need to rethink privacy policy and rules applied to research. Alternatively, journals may need to alter their publication standards to prevent future data mining of human subject data.

For ensuring data integrity in publications the analysis of the SVG code for publication graphics can ensure that figures faithfully represent the text descriptions of the publication. My detection of graphing errors in publications (wrong shapes, colours or missing data) using this method suggests that automated methods of figure verification could and should likely be used by journals during the preparation of manuscripts for publication.

## Methods

PDF documents were opened with system viewer *Preview* (version 7.0 on Mac OSX 10.9.5), the page of interest was coppied and saved as a separate PDF file. PDF was edited in *Inkscape* (Version 0.91 on Mac OSX 10.9.5) to remove non-data point information, tick marks, axis labels, text areas and shading. Then the image was saved as a scalable vector graphics file. A text parser was coded in Java to identify path objects representing data points and to extract the X and Y coordinate data, number the data point and patient. Additionally, the parser recognised different colours or shapes of the path objects to denote different patient classes. The parser saved a table as a tab-delimited text file. The text file was opened into R (version 3.2) for graphing using the library *ggplots2*(9) and axis data ranges were rescaled using the library scales(10). All code used in this analysis is in Supplementary file 1 displayed with produced tables and figures

## Supporting Information

**S1 file.** An R notebook PDF of the data code and workflow, including the Java code used to parse the SVG files.

**S2 file.** The SVG file used to extract the data and generate the plot in figure 1 D and E.

**S3 file.** The SVG file used to extract the data and generate the plot in figure 1 G.

## Acknowledgements

The author thanks UofT Coders group and Frances Wong for feedback.

## Author contribution

BC conceived the project, performed the analysis and wrote the manuscript. BC is supported by a Canada Research Chair in Maternal Fetal Health.

## References

1. Lecun Y, Bengio Y, Hinton G. Deep learning. Nature. 2015;521(7553):436–44.

2. Fienberg SE. SHARING STATISTICAL DATA IN THE BIOMEDICAL AND HEALTH SCIENCES:Ethical, Institutional, Legal, and Professional Dimensions. Annu Rev Public Health. 1994;15:1–18.

3. McGuire AL, Caulfield T, Cho MK. Science and Society - Research ethics and the challenge of whole-genome sequencing. Nat Rev Genet. 2008;9:152–6.

4. Editorial. Show the dots in plots. Nat Biomed Eng. 2017;1(5):2017.

5. Editorial. Kick the bar chart habit. Nat Methods. 2014;11(2):113.

6. Marusic A, Wager E, Utrobicic A, Hr R, Sambunjak D, Marusic A, et al. Interventions to prevent misconduct and promote integrity in research and publication. Cochrane Database Syst Rev. 2016;(4).

7. Levine RJ, Maynard SE, Qian C, Lim K-H, England LJ, Yu KF, et al. Circulating angiogenic factors and the risk of preeclampsia. N Engl J Med [Internet]. 2004 Feb 12 [cited 2011 Apr 20];350(7):672–83. Available from: http://www.ncbi.nlm.nih.gov/pubmed/14764923

8. Roberts JM, Bell MJ. If we know so much about preeclampsia, why haven’t we cured the disease? J Reprod Immunol [Internet]. Elsevier Ireland Ltd; 2013 Jul 24 [cited 2013 Aug 12];99(1–2):1–9. Available from: http://www.ncbi.nlm.nih.gov/pubmed/23890710

9. Wickham H. ggplot2: Elegant Graphics for Data Analysis [Internet]. Springer-Verlag New York; 2009. Available from: http://ggplot2.org

10. Wickham H. scales: Scale Functions for Visualization [Internet]. 2017. Available from: https://cran.r-project.org/package=scales

